# A framework for closed-loop neurofeedback for real-time EEG decoding

**DOI:** 10.1101/834713

**Authors:** Greta Tuckute, Sofie Therese Hansen, Troels Wesenberg Kjaer, Lars Kai Hansen

## Abstract

Neurofeedback based on real-time brain imaging allows for targeted training of brain activity with demonstrated clinical applications. A rapid technical development of electroencephalography (EEG)-based systems and an increasing interest in cognitive training has lead to a call for accessible and adaptable software frameworks. Here, we present and outline the core components of a novel open-source neurofeedback framework based on scalp EEG signals for real-time neuroimaging, state classification and closed-loop feedback.

The software framework includes real-time signal preprocessing, adaptive artifact rejection, and cognitive state classification from scalp EEG. The framework is implemented using exclusively Python source code to allow for diverse functionality, high modularity, and easy extendibility of software development depending on the experimenter’s needs.

As a proof of concept, we demonstrate the functionality of our new software framework by implementing an attention training paradigm using a consumer-grade, dry-electrode EEG system. Twenty-two participants were trained on a single neurofeedback session with behavioral pre- and post-training sessions within three consecutive days. We demonstrate a mean decoding error rate of 34.3% (chance=50%) of subjective attentional states. Hence, cognitive states were decoded in real-time by continuously updating classification models on recently recorded EEG data without the need for any EEG recordings prior to the neurofeedback session.

The proposed software framework allows a wide range of users to actively engage in the development of novel neurofeedback tools to accelerate improvements in neurofeedback as a translational and therapeutic tool.

## 1. Introduction

Neurofeedback uses real-time modulation of brain activity to regulate or enhance brain function and behavioral performance (for reviews, see [1, 2, 3, 4]). A closed-loop feedback approach refers to a continuous monitoring of brain activity which forms the basis for a signal that is sent back to the user of the system in real time (for reviews, see [5, 4]). Hence, closed-loop neurofeedback allows for interference with brain activity in real-time and thereby enables investigations of how brain network dynamics contribute to task performance in a causal manner, making it a powerful neuroscientific tool (for reviews, see [6, 7]). Moreover, based on recent methodological and technical advances in brain-computer interfaces (BCI), there is an increasing interest in using neurofeedback for cognitive training, especially if employed in accessible and user-friendly modalities ([8, 9], for recent reviews, see [10, 11, 12]).

The premise of using neurofeedback for cognitive training relies on self-regulation to modulate specific neurophysiological patterns. This notion was first established in the late 1950’s with experiments showing that humans were able to self-control electroencephalography (EEG) signals in real-time [13]. Learning effects, as quantified from changes in neural dynamics in response to neurofeedback, have since then been demonstrated [14, 15, 16] (rodent animal models). The specificity of neural training has been demonstrated using single-neuron activity in both non-human primates [17, 18, 19, 20, 21] and humans (e.g. [22, 23], for review, see [24]), thus providing evidence for the applicability of neurofeedback for cognitive training.

The most frequently used brain imaging modalities for neurofeedback are EEG and functional magnetic resonance imaging (fMRI) in combination with visual feedback to facilitate self-regulation of the putative brain regions that cause a specific behavior or pathology. EEG represents a low-cost, robust, and potentially mobile measurement modality, and its high temporal resolution makes it ideal for real-time neurofeedback applications. Neurofeedback in fMRI enables participants to regulate their brain activity with high spatial resolution and, in extension, provides the experimenter with information of the brain regions implicated in the task at hand.

Most published studies demonstrating the efficacy of neurofeedback in clinical settings are EEG-based. Particularly, neurofeedback for treatment of Attention Deficit Hyperactivity Disorder (ADHD) in children has been successful (for reviews, see [25, 26, 27, 28, 29]). EEG-based neurofeedback has been demonstrated to have an effect on cognitive abilities after traumatic brain injury (TBI) (for reviews, see [30, 31]). Additionally, neurofeedback and BCIs have been shown to successfully improve physical performance and neurological rehabilitation of movement disorders (for reviews, see [32, 33, 34, 35]) and for decreasing seizure incidence in epilepsy (for reviews, see [36, 37]). However, it is important to note that abovementioned meta-studies and reviews highlight the need for standardization of neurofeedback protocols and implementation, homogeneity of evaluation metrics and double-blinded study designs to yield further feasible clinical applications in the near future.

Regarding the use of neurofeedback in healthy individuals, also denoted as the “optimal” or “peak performance” field [38], there is evidence in support of cognitive performance enhancement (see three part review [1, 39, 40]). Another field of research which holds great potential and is still relatively unexplored, is the use of neurofeedback for mitigating cognitive impairment in the aging population (for review, see [3]).

While neurofeedback using fMRI has been validated in scientific settings to regulate activity in target brain regions with a high spatial resolution (e.g. [41, 42, 43, 44, 45, 46, 47], for reviews, see: [48, 49]), neurofeedback in fMRI has not reached the same potential in clinical settings as EEG-based systems, potentially due to its extensive and expensive acquisition procedures. Nonetheless, multiple studies investigate the efficacy of real-time fMRI for translational use. For example, a rapidly growing field is the use of neurofeedback for treatment of psychiatric disease. Three recent reviews [50, 51, 52] find evidence for regulation of brain regions related to emotional control using real-time fMRI, for instance for alleviating symptoms of depression or post-traumatic stress disorder (PTSD). These recent advances in neurofeedback underlines its potential for cognitive training in both healthy individuals and patients, especially if implemented in more accessible and adaptable modalities such as EEG.

Thus, developing easy-to-use software tools for EEG-based systems can accelerate investigations and implementation of neurofeedback paradigms for both cognitive training and investigations of brain-behavior relationships.

In this study, we present and evaluate a robust, closed-loop neurofeedback software framework for real-time EEG neuroimaging, cognitive state classification and feedback.

The currently available software for real-time EEG neurofeedback (for research) include BCI2000^1^ [53] (closed-source C++ software with possible integration with external programs), OpenVIBE^2^ [54] (open-source C++ software with possible integration with external programs), Brainstorm^3^ [55] (open-source Java/MATLAB software), BCILAB^4^ [56] (open-source MATLAB toolbox extension for EEGLAB software [56]), FieldTrip^5^ [57] (open-source MATLAB software), and NFBLab^6^ [58] (open-source XML/Python-based software).

In addition to abovementioned software framework, several commercial clinical software platforms exist, such as BrainMaster^7^, Myndlift^8^, BioGraph Infiniti^9^, Cygnet^10^, NeuroPype^11^, and several others. However, these software solutions lack flexibility, the possibility of adapting paradigms, and are closed-source (with varying price ranges).

Many of these software packages are written in non-interpreted programming languages that require more advanced programming skills compared to interpreted languages such as MATLAB or Python. In terms of interpreted programming languages, Python is rising in popularity as an alternative to MATLAB due to its quickly evolving open source libraries [59] and is free as opposed to MATLAB. NFBLab provides integration with Python, but to our best knowledge, the availability of EEG-based neurofeedback software packages is limited.

Therefore, the aim of this study is to provide a Python-based software framework for real-time decoding and neurofeedback based on cognitive states as measured from EEG. We prioritized the following:

1. Python source code for a high degree of modularity and transparency.
2. Fully automated, real-time implementation without the need for any manual control before or throughout the neurofeedback session.
3. Feasibility of using consumer-grade, portable EEG acquisition equipment with dry electrodes.
4. No need for any preliminary EEG recordings prior to the neurofeedback session.
5. System robustness that allows for decoding of cognitive states across intra- and inter-individidual variability.

In addition, we implement an attention training paradigm designed in fMRI [60] as a proof of concept of the framework.

In this technical note, we first provide a brief overview of major components of neurofeedback systems for cognitive state classification. Then we introduce a neurofeedback software framework for decoding of subjective attentional states using EEG, based on an attention training paradigm designed by deBettencourt and colleages [60]. Finally, we evaluate the performance of our software framework on twenty-two participants in a double-blinded design.

## 2. Methods

### 2.1. Neurofeedback System Components

In this section, we provide a brief overview of the core components of neurofeedback systems.

Principally, a neurofeedback system consists of five main elements [61]:

1. **Acquisition of brain signal data:** Measurements of brain activity can be obtained through several modalities which fall into two major categories (i) Invasive recording technologies such as electrocorticography (ECoG) or multi-electrode intracranial implants, and (ii) Non-invasive technologies such as functional magnetic resonance imaging (fMRI), EEG, magnetoencephalography (MEG) and functional near-infrared spectroscopy (fNIRS) (see reviews by [2, 62]). While non-invasive imaging modalities might lack either temporal/spatial resolution and exhibit low signal-to-noise ratio (SNR), the main advantage of non-invasive imaging (besides the noninvasiveness *per se*) is acquisition of data across the entire brain. Hence it is possible to image entire functional networks of regions involved in specific functions using approaches such as multivariate pattern analysis (MVPA) [63] (e.g. in terms of visual and semantic processing: [64, 65, 66]). MVPA methods have been used with success in fMRI neurofeedback studies [60, 67, 68, 69].
2. **Real-time data preprocessing:** Depending on the brain imaging modality, different data preprocessing steps are necessary. However, the major signal processing challenge across modalities is the detection and rejection of artifacts of either technical or physiological origin in order to acquire measurements of brain, and not artifactual, signals. Most common artifacts in EEG recordings arise from other electrical equipment, changing electrode impedance and eye/muscle movements (for reviews, see [70, 61]). Approaches for noise rejection include filtering, linear regression, and source decomposition methods (for review, see [71]). Importantly, the acquisition and preprocessing of data have to occur online and without significant latency.
3. **Feature extraction and classification:** Selecting, extracting and classifying brain signals of interest is the core of a neurofeedback system. The characteristics of interest (features) extracted from the brain recordings have to represent the brain patterns that one wants to modulate and provide feedback based upon. In EEG, these features are often spectral bands of EEG signals [72, 73, 74, 75, 76] or event related potentials (ERP) components of the signal [77, 78, 79]. Over the past 20 years, the number of journal articles about machine learning in neuroscience have grown continuously [80]. In terms of neurofeedback systems, machine learning methods allow for extraction, decoding, and thus training of increasingly complex features from noisy brain recordings (for reviews, see [81, 61, 80]).
4. **Feedback signal generation:** The extracted and decoded features of the brain signal have to be converted into a stimulus that can be continuously presented to the user of the system. This dynamic generation of a feedback signal based on an individual’s brain state is referred to as a closed-loop approach (for reviews, see [5, 4]). The feedback component can consist of sensory inputs (e.g. visual, auditory or haptic) or brain stimulation such as transcranial current brain stimulation (for reviews, see [82, 83, 84]).
5. **Adaptive user/learner:** To close the loop, the feedback is transmitted back to the user, or in case of brain stimulation, to a stimulator device. In this manner, output from the brain influences the feedback, but the input to the brain arises from the complex interactions with the feedback signal within the system, thus creating a “behavior in the loop” paradigm [5].

### 2.2. Proposed Neurofeedback Framework

We adapt the core components as described in section 2.1 to a visual sustained attention training paradigm initially designed by [60] in fMRI. In this paradigm, participants were asked to attend to a primed image category (faces or scenes) within a sequence of composite images in a block-wise structure (Fig. 1a). The top-down attentional states were decoded in real-time and used to provide a continuous visual feedback signal customized to each participant in a closed-loop manner.

**Figure 1:**
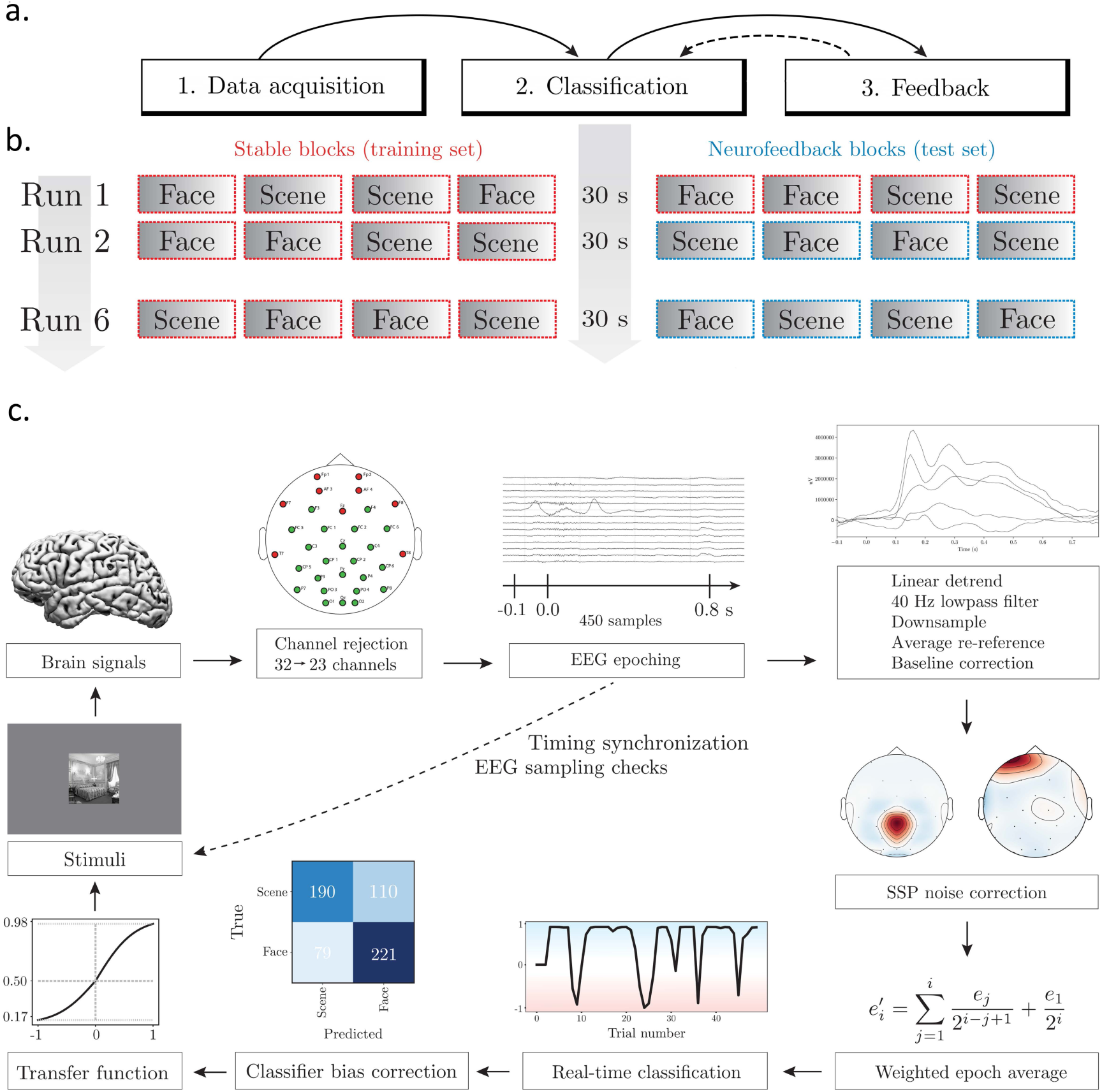
**a.** Overview of the system states. The system operates in three different states: (1) Data acquisition, (2) Classification, (3) Feedback. Data acquired during state 1 is fed into the classification state, which is then used to provide real-time feedback in state 3 (solid arrows). Data acquired during the feedback state is also used to improve the robustness of the classification models (dashed arrow). **b.** Experimental structure of the implemented paradigm. Participants completed six task runs consisting of eight blocks each. Every block (graded box) consisted of 50 trials displayed successively for one second each. The blocks with a red, dashed outline are denoted as ‘stable’ blocks, and the blue blocks are denoted as ‘feedback’ blocks. **c.** Overview of the components necessary for providing closed-loop feedback on a trial-to-trial basis.

A supplementary video of the system is available (section 6).

The system operates in three different states: (1) Data acquisition, (2) Classification, (3) Feedback (Fig. 1a). For the acquisition state, data is solely recorded and no feedback is provided. The data recorded during this state is used for training a model capable of classifying cognitive state. During the classification state, the recorded data is first used to identify artifactual data components for each individual participant. Second, a classification model is trained on the most recently recorded data. The classification step can take place during breaks in the experimental paradigm. The allocated time for the classification step depends on the model and data complexity, as well as on computing resources. Ultimately, the feedback state updates the stimulus on a trial-to-trial basis. The components necessary for providing closed-loop feedback on this trial-to-trial basis are illustrated in Fig. 1b. The data acquired during the feedback state is also exploited during the subsequent classification state (dashed line in Fig. 1a).

#### 2.2.1. EEG Acquisition Equipment

A consumer-grade, portable EEG equipment, Enobio (Neuroelectrics) with 32 dry-electrode channels, was used for data acquisition. The EEG was electrically referenced using a CMS/DRL ear clip. The system recorded EEG data with a sampling rate of 500 Hz with EEG electrodes positioned according to the 10–20 system. The data was transmitted to a Lenovo Legion Y520 laptop via a USB cable. Visual stimuli were presented on a dual monitor. Lab Streaming Layer^12^ (LSL) was used for streaming and synchronizing EEG data with feedback signal generation. LSL supports short latency data exchange and buffering for the most common EEG/MEG devices.

#### 2.2.2. Real-Time EEG Preprocessing

EEG epochs of 900 ms (100 ms pre-stimulus onset, 800 ms post-stimulus onset) were extracted for each image trial and preprocessed epoch-wise in real-time. The EEG signal was linearly detrended and lowpass filtered with a cut-off frequency of 40 Hz using a Finite Impulse Response (FIR) filter with zero-double phase. The EEG signal was downsampled from 500 Hz to 100 Hz and referenced to an average of the 23 preselected channels (see below). Baseline correction was performed based on 100 ms of the pre-stimulus signal. EEG epochs were z-scored such that each trial had a mean of 0 and a standard deviation of 1.

Feedback estimates were based on pre-selected spatial areas (EEG channels). Based on pilot data and the recommendations of [85], nine frontal and temporal channels (Fp1, Fp2, Fz, AF3, AF4, T7, T8, F7, F8) were rejected in all real-time analyses. These channels contained a high level of muscle and eye artifacts. Besides these nine channels, no channels or epochs were rejected real-time. Besides the option to pre-select/reject specific EEG channels for all real-time analyses, the software framework provides an option for dynamically rejecting channels if they contain signal either above or below given thresholds, thus handling faulty electrodes. In that case, any rejected channel would be re-estimated based on interpolation of its neighbors. No epochs were rejected during preprocessing. The signal was artifact corrected using signal-space projection (SSP) (section 2.2.3).

The preprocessing schemes were largely based on implementations provided by MNE software [86].

#### 2.2.3. Real-Time Artifact Rejection: Signal-Space Projection

We employed a data decomposition approach using signal-space projection (SSP) for detecting and rejecting artifacts in real-time. SSP relies on the fact that the electric field distributions generated by the sources in the brain have spatial distributions sufficiently different from those generated by external noise sources. SSP does not require additional reference sensors to record the disturbance fields [87, 88], which makes it suited for real-time implementation.

For the mathematical modeling of the aquired signals we use two representations. We use a matrix representation where the multi-channel signal is kept in a channel-by-time matrix **X**. For a multi-epoch signal we concatenate the channel time series. When convenient, we also use a vector representation **x** in which we concatenate the time series of the retained channels, typically for a single epoch.

In essence, SSP decomposes the noisy data into components by a singular value decomposition (SVD) of the signal. It is important to note that these components might not be statistically independent, and therefore there is a risk that brain signals of interest and artifacts may be reduced in the de-noising process [87].

An SVD was performed on the matrix of concatenated epochs of noisy EEG events (the training set), **X**_training_ [88]:

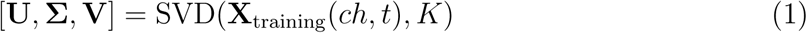

Based on the SVD, the number of principal components (PC) was chosen such that each PC independently explained at least 10% variance of the total variance in **X**_training_. Suppose there are *K* such PCs, then the corresponding singular vectors in **U** are projected out of future noisy data (the test set-as subject to neurofeedback) to obtain a noise-corrected signal, **X**_clean test_:

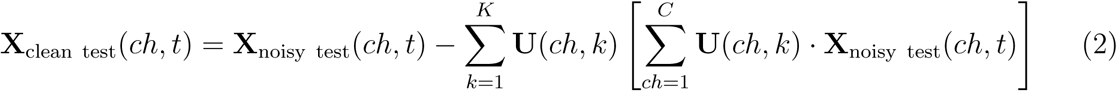

In the current framework, components explaining more than 10% variance were identified based on ‘stable’ blocks (training set) and subsequently rejected for each EEG epoch in the ‘feedback’ blocks (test set). The SSP projectors were re-estimated after each set of ‘stable’ blocks to ensure an updated estimate of the signal composition throughout the experiment.

The real-time SSP implementation was based on an MNE framework [86].

#### 2.2.4. Real-Time EEG Classification

The classification task consisted of categorizing the top-down attentional states towards faces and scenes, i.e. a binary classification task.

The logistic regression model models the posterior probabilities of two classes based on EEG features, **x**. The probability of assigning an observation **x**_*i*_ to class *y* = 1 is,

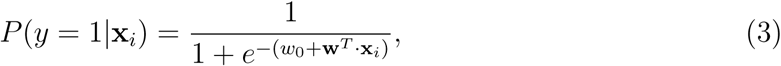

where the parameter set (*w*_0_, **w**) is estimated on a training set.

The implemented model was a logistic regression classifier with L1 norm regularization from the scikit-learn toolbox [89] using the ‘saga’ solver [90] with a *C* regularization of 1 (default setting). Due to the accessibility of the scikit-learn toolbox, the framework provides easy adaptability to various classification models to suit the experimenter’s needs.

In brief, participants completed six task runs consisting of eight blocks each. Every block consisted of 50 trials (1 second each). The very first task run consisted of eight blocks used solely for data acquisition (‘stable’ blocks), while the remaining task runs consisted of four ‘stable’ blocks, followed by four ‘feedback’ blocks (see Fig. 1a and section 2.2.8 for description of the experimental structure).

For each task run besides the very first one, a classifier was trained in the 30 s break between ‘stable’ and ‘feedback’ blocks. For the first task run, the classifier was trained on the first 600 ‘stable’ EEG trials (twelve blocks). For the following task runs, the classifier was trained on the 600 most recent ‘stable’ EEG trials and the 200 most recent ‘feedback’ trials (illustrated as dashed line in Fig. 1a) (sixteen blocks in total). In this manner, the training data for the classifier were continuously updated throughout the experimental session to include the most recent sixteen blocks corresponding to 800 trials.

For artifact rejection, SSP projectors were computed based on the twelve stable blocks in the training data (see section 2.2.3). Following artifact rejection, two consecutive epochs were averaged over a moving window across all training epochs. Finally, a classifier was trained on the artifact-corrected and averaged data.

The trained classifier was tested in real-time on EEG data epochs recorded during the ‘feedback’ blocks. A test epoch was preprocessed and corrected using the SSP projectors computed based on the training data as described above.

The preprocessing, artifact rejection and decoding of an epoch, *e*, was performed during the last 200 ms of the current experimental trial. Following the real-time data processing of the current epoch, each test epoch was averaged using a type of exponentially weighted moving average. Thus, the current epoch, *e*, was updated:

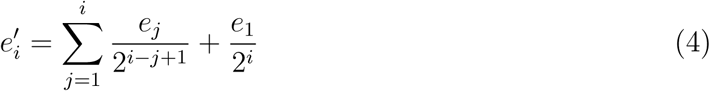

Hence, the classifier was tested on a weighted moving average of the most recent trials with 50% weight on the current trial, and exponentially decreasing weight on earlier trials.

The implemented logistic model estimated a prediction probability, *p*_*c*_, i.e. the confidence of the classifier that a trial belonged to the task-relevant category. The prediction probability of the classifier was transformed into a classifier output for each test epoch, following the framework in [60]. The classifier output was given as:

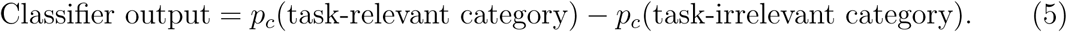

The classifier output ranged from −1 to 1, with values above 0 indicating task-relevant category information decoded in the participant‘s brain and values below 0 indicating task-irrelevant decoded information. The classifier output was fed into a transfer function and used to generate a feedback signal to the participant, as described in section 2.2.5.

Based on pilot data, it was observed that the trained classifier was frequently biased towards one of the categories (often the face category). The prediction probabilities of the feedback epochs were therefore adjusted using an offset calculated based on the training data. The offset for the first ‘feedback’ run was computed on the training data in a 3-fold cross-validation procedure. For the remaining ‘feedback’ runs, the offset was based on the bias of the four most recent ‘feedback’ blocks (200 trials). The bias offset was limited to an absolute value of 0.125.

#### 2.2.5 Real-Time Feedback Generation

The classifier output eq. (5) computed in real-time was mapped to a relevant neuro-feedback stimulus value following the approach proposed in [60]. In particular, the classifier output was mapped to a visual feedback stimulus, i.e. proportion of task-relevant category in the composite image (*α*), using a shifted and scaled sigmoid function as shown in Fig. 2. Based on pilot studies, the mapping was designed to map the classifier output to a sensitive range of values for updating stimuli in a closed-loop manner, yet avoid saturation. A classifier output of −1, i.e. when the classifier assigned maximum prediction probability to the task-relevant category would yield the task-relevant image to be 98% visible. Conversely, a classifier output of 1, i.e. when the classifier assigned maximum prediction probability to the task-irrelevant category would yield the task-relevant image to be 17% visible. Thus, the output of the function ranged from 0.17 to 0.98. The minimum value of 0.17 ensured that the task-relevant category was always visible to some degree in the composite image, giving the participant a chance to recover from an attention lapse.

**Figure 2:**
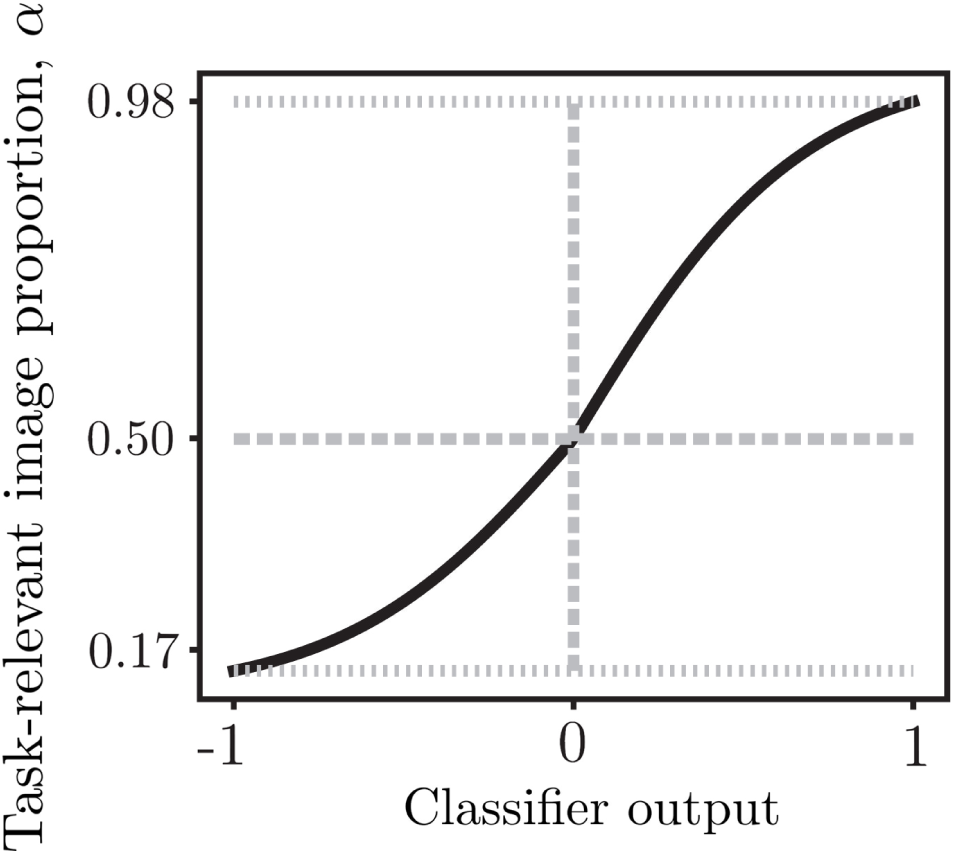
Sigmoid transfer function for mapping the classifier output to a task-relevant image proportion (interpolation factor *α*) for visual feedback generation. The sigmoidal function was asymmetric with an inflection point of 0.6.

Since (i) Sustained attention has been shown to fluctuate slower than on a trial-to-trial basis [91, 92, 93], and (ii) The classifier output contains misclassifications that can be described as high-frequency noise, the three most recent *α* values from the transfer function were averaged to provide a mixture proportion for updating image trials.

#### 2.2.6. Participants

Twenty-two healthy adults (10 female, 22 right-handed, mean age = 22.9, age range = 20-26) with normal or corrected-to-normal vision participated in the study for monetary compensation. This included an EEG neurofeedback group and a control group receiving sham neurofeedback. Each participant in the control group was matched randomly to a participant in the neurofeedback group based on gender. The matched pairs were presented to identical stimuli on all three days, and control participants thus received yoked feedback based on the decoded neural response profile of their matched neurofeedback participant. The yoking ensured that control participants were exposed to the same variations in image mixture and task difficulty. No participants were excluded from the study. The study was performed with automated randomization and group assignment to achieve a double-blinded experimental design. The first few participants were necessarily assigned to the neurofeedback group to provide matched, yoked feedback to the control group, and similarly the last few participants were assigned to the control group. Thus, the experimenter was blind to group assignment in 16 out of 22 participants (72.7%). All participants received the same scripted instructions.

All participants provided written informed consent to a protocol approved by The Institutional Ethical Review Board, University of Copenhagen, Department of Psychology (approval number: IP-IRB / 26112018) following the 1975 Helsinki Declaration as revised in 2008.

#### 2.2.7. Experimental Stimuli

Images consisted of grayscale photographs of female/male faces and indoor/outdoor scenes that were combined into composite stimuli by interpolating two images using a variable interpolation factor, (*α*). All image trials were generated from an interpolation of two random images, choosing from 1000 images for each subcategory, i.e. a total of 4000 unique images. The face images were from the FERET database^13^ [94] and scene images were from the SUN database^14^ [95]. All images were resized and cropped to 175 x 175 pixels. To avoid the influence of low-level visual features, we ensured that all images had a similar brightness level. The pixel values of the images were linearly transformed such that the following two groups of values each accounted for 10% of the pixel values: 0-46 and 210-255. For the face images, the quantiles were estimated based on the center of the images, such that the “dead space” around the face was not included. A white fixation cross was superimposed on the images and presented during breaks, except when text instructions were displayed. The images were presented against a mid-gray background. Participants were seated approximately 57 cm from the computer monitor, corresponding to a 5 degree visual angle.

#### 2.2.8. Experimental Neurofeedback Paradigm

An attention training paradigm was implemented, as designed in [60]. Participants were asked to pay attention to a subcategory of *faces* or *scenes* in a sequence of composite images (e.g. a mixture of 50% of an image of a *female* face and 50% of an image of an *indoor* scene). The neurofeedback element consisted of updating the image proportion (interpolation factor, *α*) of the composite images (e.g. a mixture of 80% of an image of a *female* face and 20% of an image of an *indoor* scene) based on the participant‘s decoded attentional level to the primed image category.

Participants underwent six task runs, where each task run contained eight blocks with 50 trials in each block (total of 2400 trials). Each block began with a text cue for 6 s that instructed participants which subcategory was the target to which they should focus their attention towards and respond to by a keypress. The text cues were *’indoor’*, *’outdoor’*, *‘female’* and *’male’*. For each block, the text cue was followed by 2 s of fixation (white cross centered on a mid-grey background) before the sequence of 50 image trials started. The image trials consisted of composite face/scene images that were displayed for 1 s each with no inter-stimulus interval (ISI).

The first task run only contained blocks with ‘stable’ trials, i.e. an equal mixture of faces/scenes for each image. Starting with the second run, the first four blocks were ‘stable’, and the following four blocks were ‘feedback’ blocks with a variable mixture proportion depending on the attentional level decoded from EEG recordings of each participant (in the neurofeedback group).

Participants were informed by text on the screen each time a new type of block was initialized. For each ‘feedback’ block, the first three image trials had equal mixture proportion to allow time for classifier decoding of presented trials. The mixture proportion for the remaining trials (47 trials) within each block were based on the real-time EEG classifier output of decoded attentional level. If participants attended well (high levels of task-relevant information decoded in their brain) the task-relevant image became easier to see, and vice versa. Thus, the feedback worked as an amplifier of participants attentional state, with the goal of making participants aware of attention fluctuations and hence improve sustained attention abilities.

#### 2.2.9. Experimental Procedure

All participants completed two behavioral sessions and one neurofeedback session on three consecutive days. The first day was a behavioral pre-training session with two runs of the sustained attention task without recording of EEG. The second day was an EEG session consisting of a single run with stable stimuli as in the pre-training session, and five runs with real-time neurofeedback. The third day was a behavioral post-training session with two runs of the attention task without recording of EEG, similar to the first day.

Participants were asked to pay attention to the primed target category, and respond to to a response inhibition task [96, 97]. 90% of the images (45 images in each block) contained the target category and required a keypress response, while 10% (5 images in each block) contained the non-target category to which responses had to be withheld.

For each task run, four of eight blocks involved attending to faces and the remaining four blocks involved attending to scenes. To avoid an unbalanced category distribution, the target categories were randomly assigned to each block with the constraint that two blocks of each target category had to be present within the first and last four blocks.

The target subcategories presented for each participant throughout the three-day experiment were held constant, i.e. if the target categories were *’indoor’* and *‘female’* for a particular participant during the pre-training session on day 1, the participant would be primed to the same categories on the following days. The category assignment for each participant was random, but counterbalanced across all participants. Since the task runs contained randomly generated target categories and composite image trials, each run was unique to avoid a habituation and recognition effect.

For behavioral sessions during the pre-training (day 1) and post-training (day 3) sessions, all participants completed two task runs of eight blocks (total of 800 trials). For these behavioral sessions, all composite image trials had an equal mixture proportion of face (50%) and scene (50%), denoted as ‘stable’ images.

During both the behavioral and EEG sessions, participants were given 30 s breaks after four blocks. The total experiment time was 17 min for the behavioral session and 52.6 min for the EEG session including breaks, text cues and fixation time.

For each of the three sessions, participants were instructed to sit relaxed in a chair with their right hand resting on the table with a finger on a keyboard for providing behavioral responses. Participants were instructed to keep their eyes focused on the fixation cross on the screen. Previous to the EEG session, participants were asked to avoid excessive movements during stimuli presentation. Participants were informed that the ‘feedback’ trials would be updated based on their attention towards the target category and not based on their keypress response. Specifically, the task-relevant image would become easier to see if they were paying attention and harder to see if their attentional level was decreasing. For the pre-training and EEG session, participants were shown short examples of respectively the behavioral experimental paradigm and neurofeedback paradigm.

### 2.3. Offline Data Analysis

#### 2.3.1. Offline EEG Classification

The ability to decode attentional states within participants was assessed by classification of ‘stable’ blocks (28 blocks) in a leave-one-run-out (LORO) cross-validation. ‘feedback’ blocks were not included in the cross-validation, since participants received different types of feedback. Similarly to the real-time pipeline, epochs in the training blocks were preprocessed epoch-wise and SSP projectors were computed and applied to the full training set. Following SSP correction, two consecutive trials were averaged over a moving window across the entire training set. For classifier testing, the SSP projectors computed based on the training set were applied to the test set, followed by a weighted moving averaging (Eq. (4)) across test epochs. To provide an estimate of classifier bias, the offset was computed based on a 3-fold cross-validation of the training data and applied to the prediction probabilities of test epochs.

For the LORO approach, all ‘stable’ blocks within a single run was in turn used as a test set once (besides the very first run which only consisted of ‘stable’ blocks. This run was always included in the training set to avoid variability in the size of test set).

The classification performance was evaluated as decoding error rate (i.e. how often the classifier predicted the wrong category).

#### 2.3.2. Sensitivity Mapping of Spatial and Temporal Features Important to Classification

We implemented a sensitivity map to visualize the EEG signatures exploited by the real-time classifiers. The employed classifiers (Eq. (3)) weigh the input features. The sensitivity map, **M**, visualizes the importance of individual features spatially, at each channel (ch), and temporally, at a relative time (*t*):

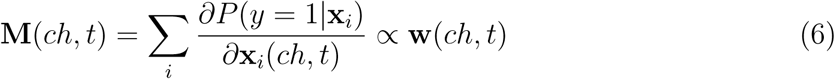

where **w** are the weights of the L1 logistic classifier. The sum on samples extends over the given training set.

To compute the uncertainty of sensitivity maps, we use a NPAIRS (nonparametric prediction, activation, influence, and reproducibility resampling) split-half resampling scheme [98]. The NPAIRS workflow randomly divides the sample in two half-sized sets. By exchangeability the squared difference of the maps for a given split is an unbiased estimator of the variance. Averaging over multiple splits (*S*) we reduce variance of the unbiased estimator. We subsequently scale the average map to produce an effect size map 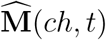. In particular, we compute the sensitivity map on a split consisting of 11 random participants:

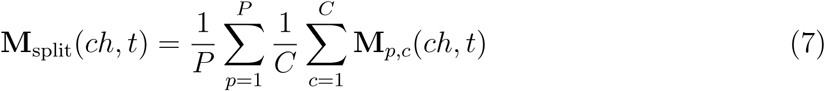

subtract and square to provide an unbiased feature-wise uncertainty measure:

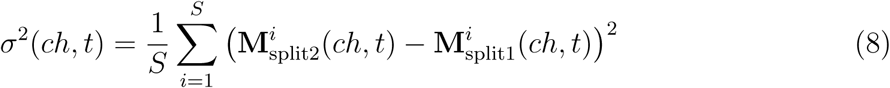

which is finally used to arrive at an effect size map:

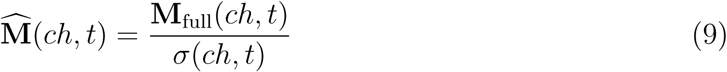

where **M**_full_(*ch, t*) is the sensitivity map estimated based on the full set of training data (all participants). The scaling was calculated based on *S* = 10, 000 resampling splits. In each split, the participants were sampled without replacement.

#### 2.3.3. Behavioral Performance Metrics

Keypress responses were collected from participants on all three experimental days. Keypress responses were extracted 150 ms to 1150 ms post-stimulus onset. In case of multiple keypresses for a single image trial, the first keypress response was used. Measures of behavioral performance were computed as error rate and a non-parametric index of sensitivity, A’ [99]:

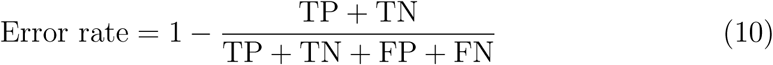

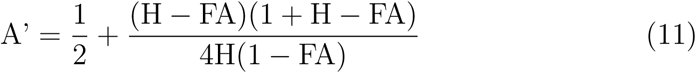

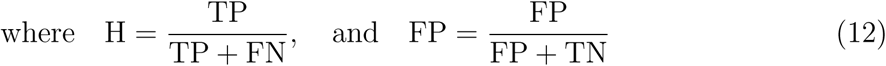

True positives (TP) were responses during non-lure trials, false negatives (FN) were rejections during non-lure trials, true negative (TN) were rejections during lure trials, and false positives (FP) were responses during lure trials.

Due to low values of error rate measures across participants, the relative error rate reduction (RERR) was computed as:

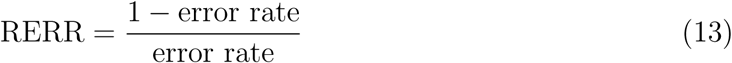

## 3. Results and Discussion

In this section, we evaluate the system performance of the software framework presented in Section 2.2. First, we report the unbiased decoding error rates of the system obtained from offline cross-validation analyses. Next, we report the decoding error rates obtained in real-time, as well as the fluctuations of real-time decoding error rates throughout the time course of the experiment. Moreover, we visualize which parts of the EEG signature were exploited by the classifiers trained in real-time to predict cognitive states. Lastly, we report and discuss the training effects in participants’ sustained attention abilities during and after the neurofeedback session.

### 3.1. Classifier Decoding Error Rate

The system‘s ability to decode attentional states was assessed by classification of ‘stable’ blocks in a leave-one-run-out (LORO) cross-validation (section 2.3.1) to provide a measure as unbiased as possible from participants’ receiving different types of feedback. The mean classifier error rate was 0.337 for neurofeedback participants (range: 0.194 - 0.446), and 0.348 for control participants (range: 0.207 - 0.467) (Fig. 3). The chance level of 0.5 was outside the 95% confidence interval (CI) for 17 out of 22 participants (95% CI computed based on five cross-validation folds).

**Figure 3:**
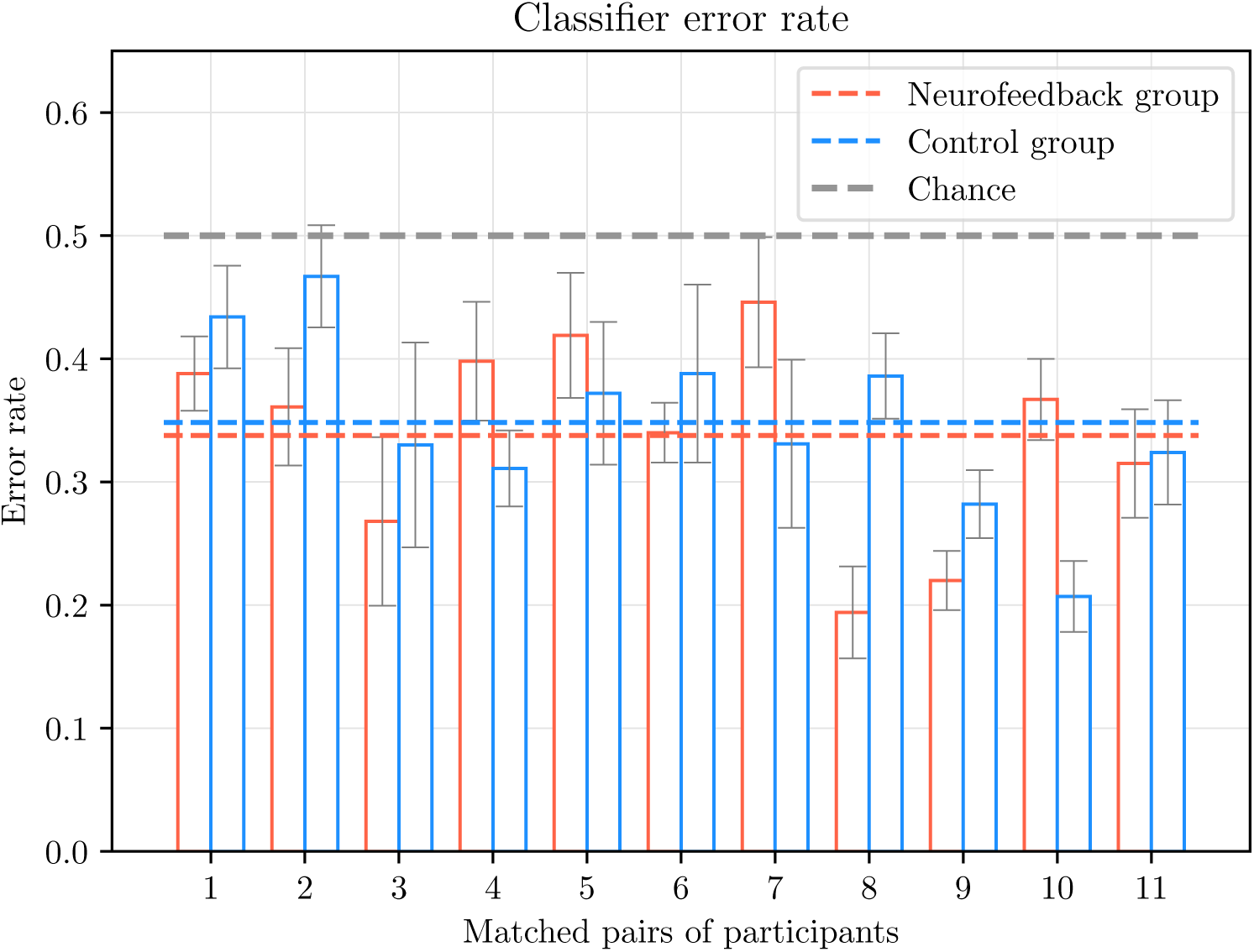
Classifier decoding error rate (estimated offline by cross-validation of ‘stable’ blocks in a leave-one-run-out approach). Red bars illustrate error rates of the participants in the neurofeedback group. The paired blue bars illustrate error rates of the matched control participant (N=22, eleven pairs of participants). Dashed lines depict mean error rate values. Error bars represent the unbiased standard error of the mean (SEM) per participant (computed as the standard deviation across five cross-validation folds without normalizing by the square root of five) [100].

Hence, we achieved a mean classifier error rate of 0.343 (N=22) using a consumer-grade, dry-electrode EEG equipment. In comparison to the original paradigm implemented in fMRI, deBettencourt and colleagues reported a mean classifier error rate of 0.22 (N=32) using MVPA methods [60]. These classifier error rates represent the ability to decode top-down attention towards faces and scenes. Participants were exposed to the exact same type of composite images for both categories (equal image mixture proportion), and thus EEG signals associated with subjective attentional states, and not solely visual responses, were decoded.

### 3.2. EEG Decoding Error Rates Obtained Real-Time

The proposed framework was capable of decoding fluctuating attentional states in real-time. This was achieved by repeatedly training classifiers on the most recently recorded EEG signals throughout the time course of the experiment (~52 min). Note that the decoding error rates obtained in real-time were influenced by the fact that participants’ received either true or sham neurofeedback and hence experienced different levels of task difficulty.

Figure 4 demonstrates the decoding error rates obtained in real-time during the ‘feedback’ blocks (20 blocks, 1000 trials per participant) during the neurofeedback session. The mean real-time decoding error rate for neurofeedback participants was 0.384 and 0.387 for control participants. For neurofeedback participants, this real-time decoding error rate was used to continuously update the stimuli, while the control participants were exposed to stimuli updated based on the decoded neural response profile from a matched neurofeedback participant. The real-time decoding error rates are not reported for the fMRI study for the same attention training paradigm [60]. The real-time error rates are lower than the classifier error rates reported in section 3.1 due to the non-stationarity of the feedback signal. Moreover, if a participant was inattentive towards the task-relevant category, a high decoding error rate would be classified as “correct”, i.e. an ability of the classifier to successfully decode lapses in subjective attentional states.

**Figure 4:**
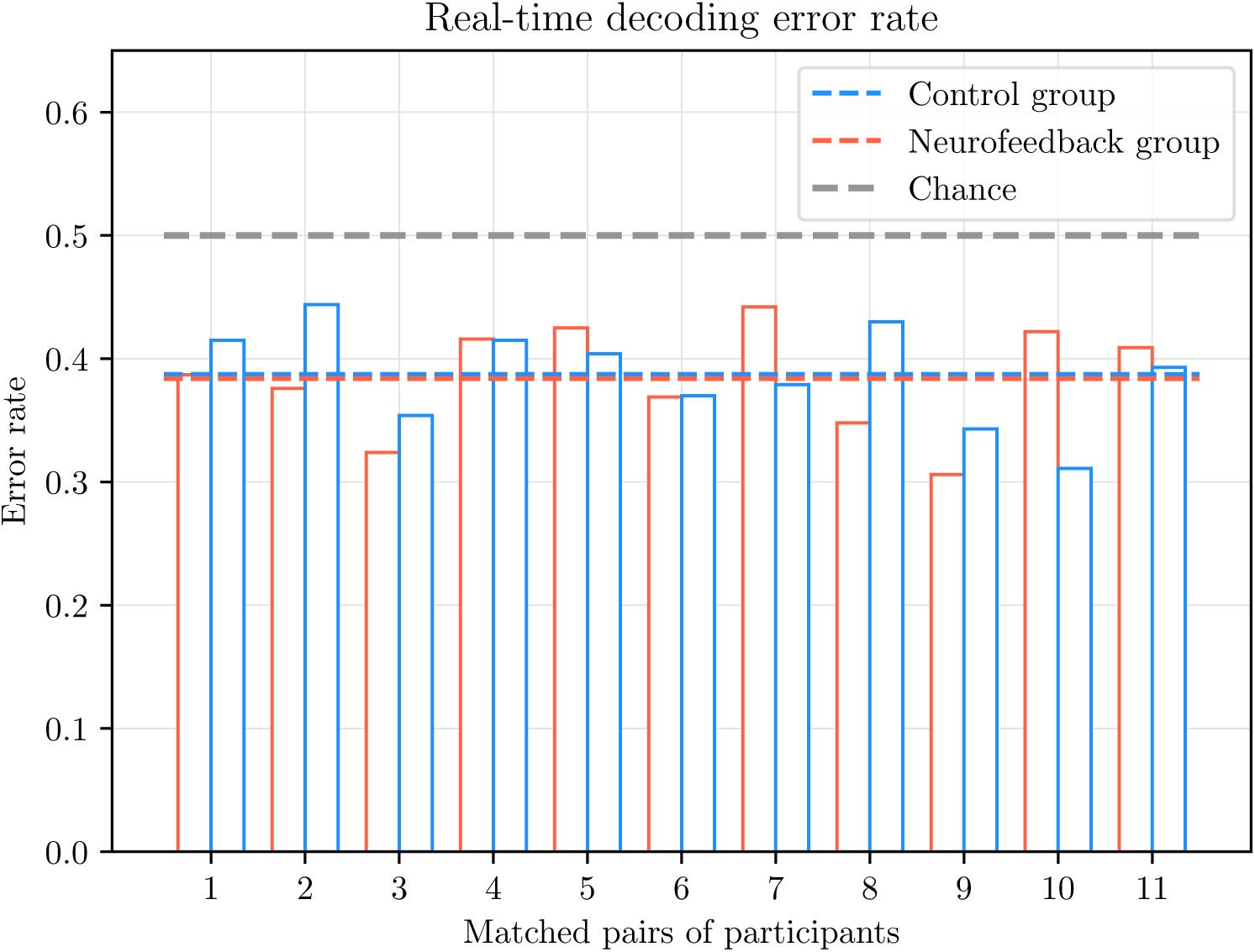
Real-time decoding error rate (obtained during ‘feedback’ blocks). Red bars illustrate error rates of the participants in the neurofeedback group. The paired blue bars illustrate error rates of the matched control participant (N=22, eleven pairs of participants). Dashed lines depict mean error rate values.

To investigate how the decoding error rate fluctuated throughout the ‘feedback’ blocks of the neurofeedback session, the mean decoding error rate across participants per ‘feedback’ run is shown in Fig. 5. The mean decoding error rates per run across all participants were as follows: 0.416, 0.38, 0.389, 0.37, 0.375. Thus, as expected the error rate had a decreasing trend across the time course of the neurofeedback session: The classifier was trained on 12 blocks for the first ‘feedback’ run (600 trials), and 16 blocks (800 trials) for the remaining ‘feedback’ runs, thus achieving lower error rates by using larger training sets.

**Figure 5:**
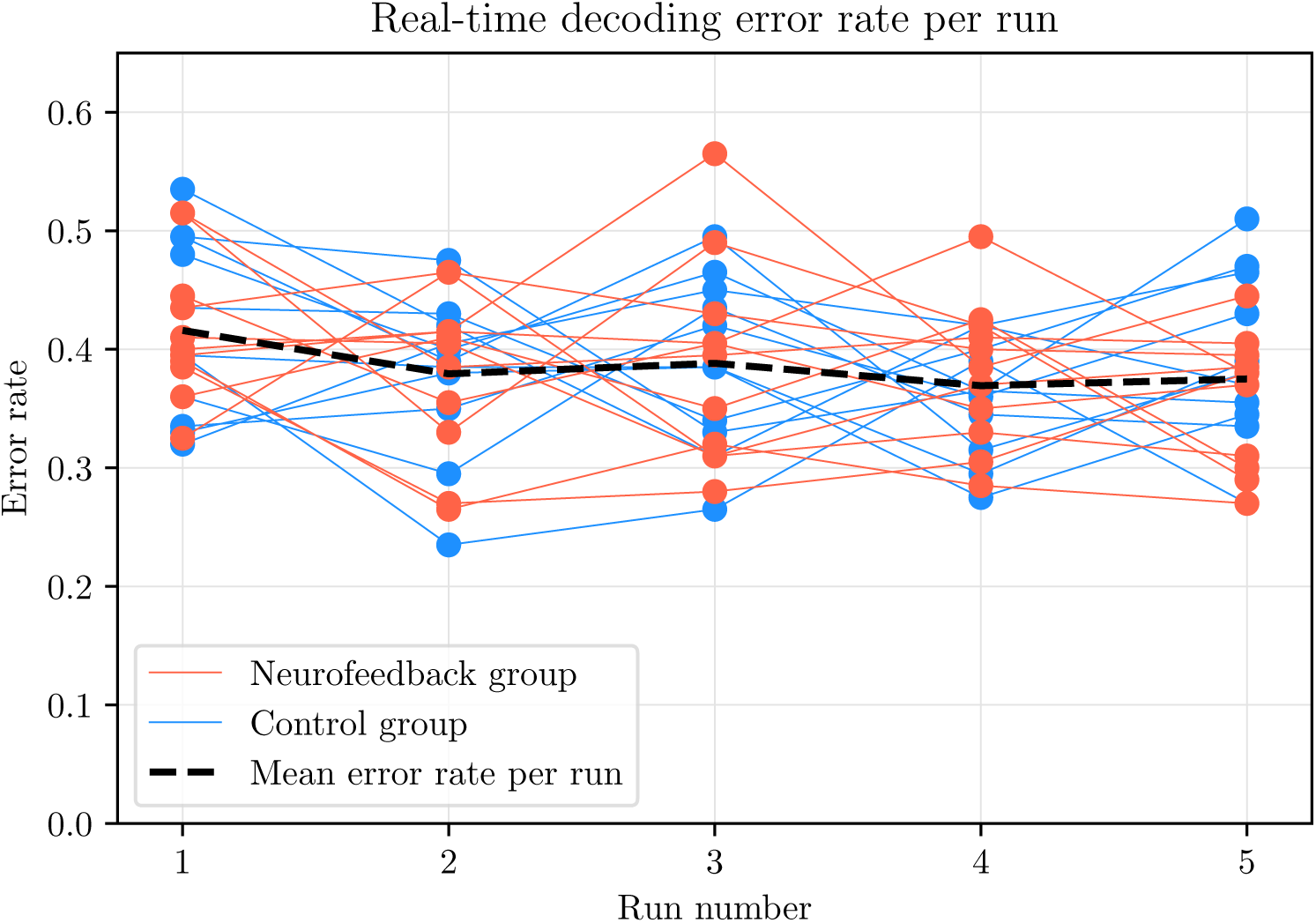
Real-time decoding error rate (obtained during ‘feedback’ blocks) per run across all participants (N=22). Black dashed line depict mean error rate values for all participants.

### 3.3. Sensitivity Map of Real-Time Classification

We investigated the temporal and spatial EEG signatures exploited by the real-time classification models. Throughout the neurofeedback session, a total of five logistic classifiers were trained on the most recently recorded EEG signals in order to appropriately decode visual top-down attentional states and provide relevant feedback. A sensitivity map was computed based on the averaged L1 logistic model weights across all real-time classifiers for all participants. For sensitivity map effect size evaluation, we implemented an NPAIRS resampling scheme [98]. In this cross-validation framework, the obtained model weights for each participant were split in two partitions of equal size (eleven participants in each partition randomly selected without replacement). This procedure was repeated 10,000 times to obtain standard error of the sensitivity maps for computing effect sizes (section 2.3.2, Fig. 6).

**Figure 6:**
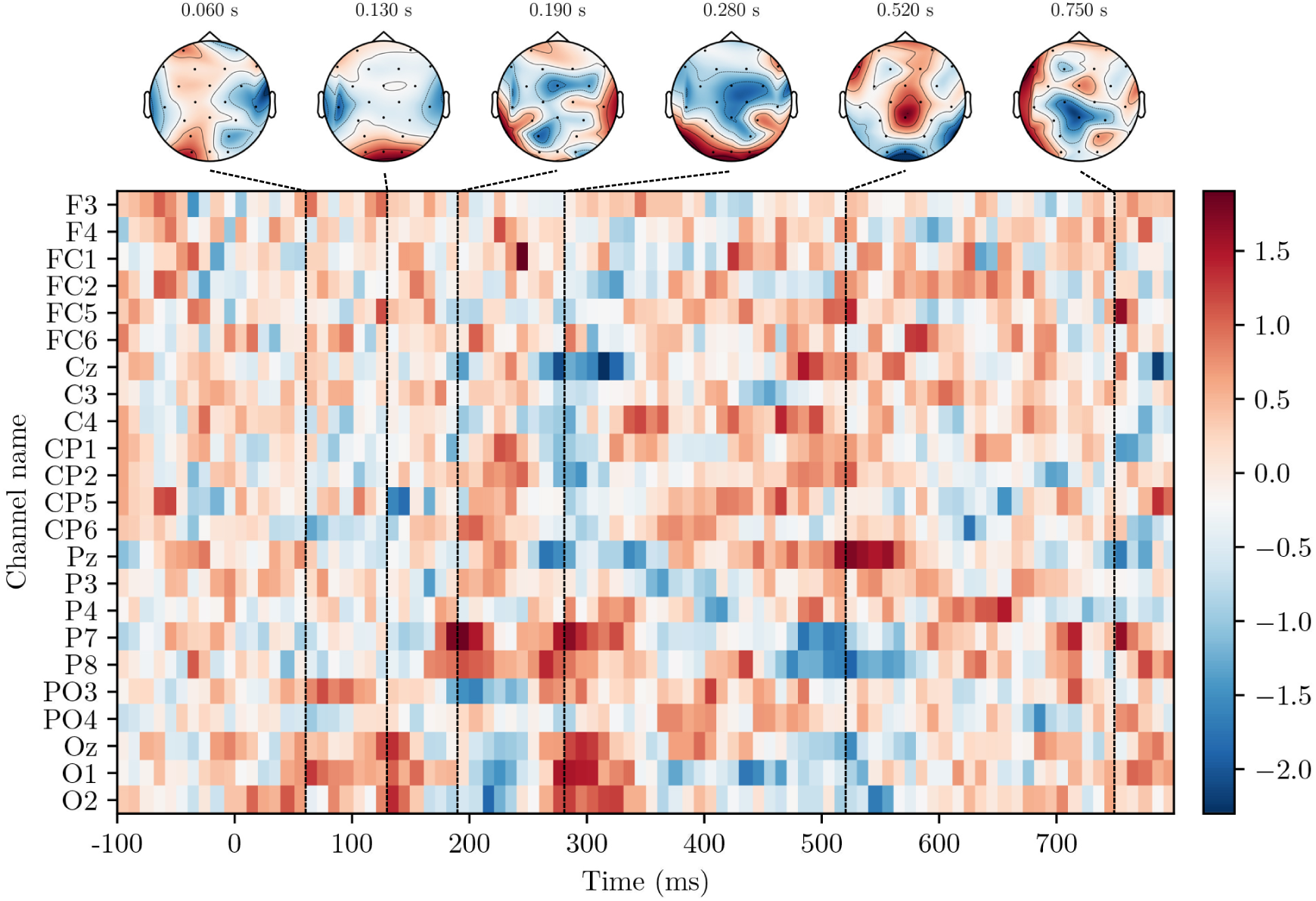
Effect sizes for the sensitivity map computed based on real-time logistic model classifiers across all participants (N=22). Effect sizes were computed based on 10,000 NPAIRS resampling splits. The x-axis displays the temporal importance of features (90 EEG samples, 900 ms), while the y-axis displays the spatial importance (23 EEG channels) relevant to the classification task performed in real-time.

As expected in decoding of visual attentional states, occipital EEG channels (O1, O2, Oz) during early visual processing (at time points around 110-160 ms) were evident on the sensitivity map (Fig. 6) consistent with the visual ERP literature (e.g. [101, 102], for review see [103]). Moreover, temporal components around 160-190 ms (P7, P8) were of importance in the classification task, potentially corresponding to the signature of face processing in the human brain (N170 component) [104].

Interestingly, the classifiers exploited later temporal components besides the early visual response which might correspond to processing of subjective visual states as proposed in previous work [78, 105, 106, 107]. Treder and Blankertz reported optimal decoding of overt attention based on early evoked potentials, and conversely optimal decoding of covert attention based on later components (P300) around parietal channels [78]. Similarly, Kasper and colleagues showed that attentional failures could be distinguished from successes during an attention task on a single-trial basis using a 400-500 ms temporal window [105].

Thiery et al. predicted covert visuospatial attention from ERPs (albeit using manually pre-defined temporal windows and occipital regions) and demonstrated that both an early time window (0-100 ms) and a later component at 410-530 ms on occipital channels were optimal for attention decoding [106]. Moreover, List and colleagues showed that the time interval 500-625 ms across occipital/parietal brain regions contained discriminative information for distinguishing locally-from globally-focused attentional states [107].

In terms of spatial features, we observe activity across not only occipital visual areas, but also distributed activity across frontocentral brain regions (Fig. 6). In fMRI, activity over frontoparietal regions has been linked to cognitive control/goal directed behavior [108] and brain regions supporting attention training process [60].

Hence, the signatures of the sensitivity map (Fig. 6) provide evidence that we were able to successfully decode subjective attentional states in real-time throughout the neurofeedback session.

**Figure 7:**
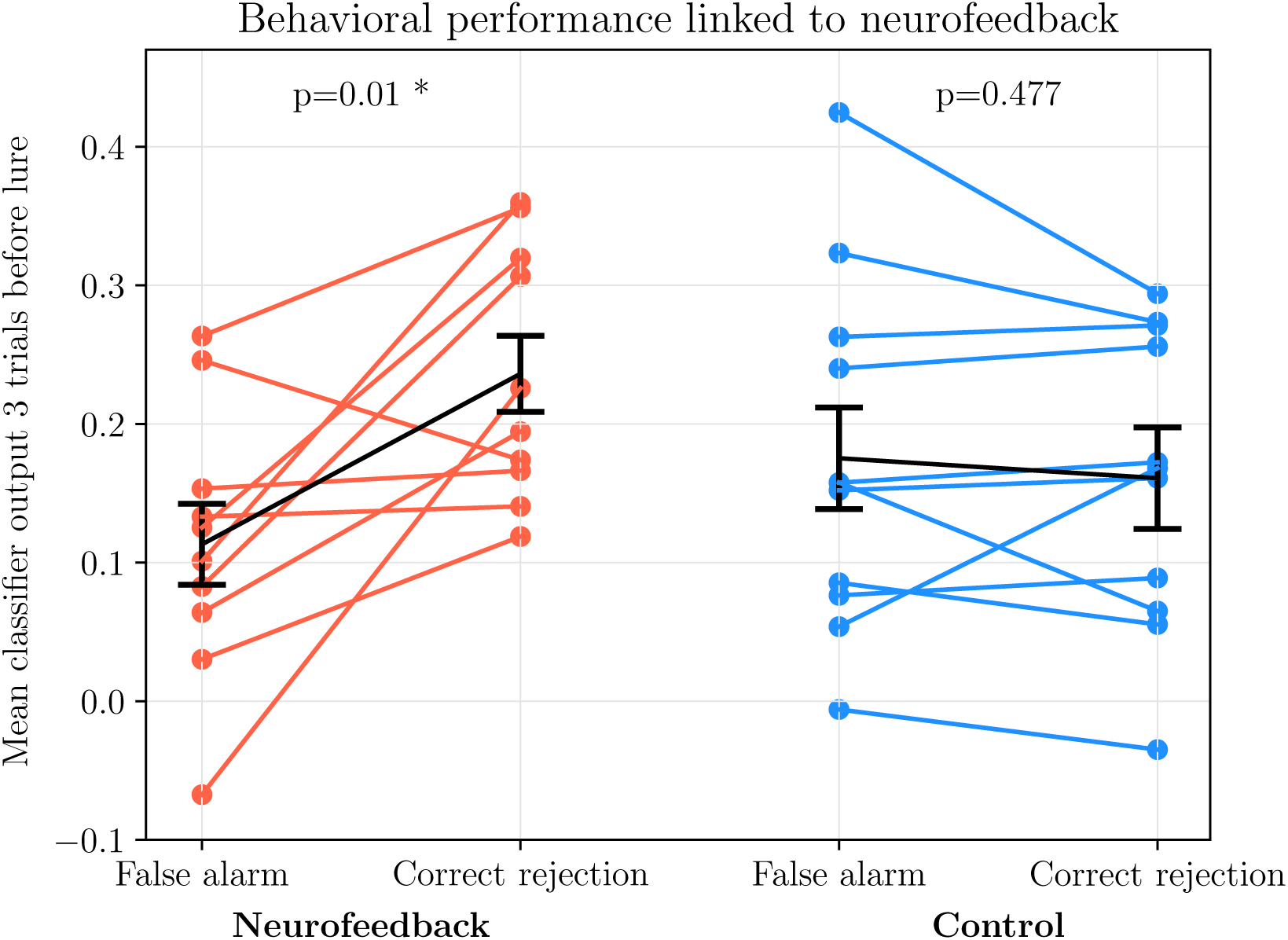
Behavioral task performance linked to decoded attentional EEG state. Error bars represent +/− 1 SEM (N=10 for neurofeedback, and N=11 for controls).

### 3.4. Training Effects of Neurofeedback

The training effect of neurofeedback was investigated both (i) During the neurofeedback session itself, and (ii) The day following the neurofeedback session.

During the neurofeedback session, we investigated whether the classifier output (i.e. the amount of task-relevant information decoded in a participant‘s brain) before a lure trial in the response inhibition task predicted whether participants correctly withheld their response or incorrectly responded to a lure trial. The classifier output was averaged over the three trials preceding the lure trial. A low classifier output indicated that the participant had a low task-relevant neural representation, and hence was inattentive. For neurofeedback participants, this low degree of task-relevant information could indeed be used to predict when a participant would make an erronous response (false alarm). Vice versa, a larger degree of task-relevant information in a neurofeedback participant‘s brain was linked to a correct response (correct rejection of a lure trial). The difference in classifier output between false alarms and correct rejections was significant for the neurofeedback group and insignificant for the control group with an interaction of p=0.7e−3. This provide evidence that we were able to control participants’ behavior using neurofeedback.

To investigate whether the single neurofeedback session provided lasting effects, we compared the behavioral performance of participants during the pre-training session (day before the neurofeedback session) and the post-training session (day after the neurofeedback session). First of all, there was no baseline difference (pre-training session) between the neurofeedback and control group in terms of A’ (p=0.9), RERR (p=0.9) or response time (p=0.8).

Second, the neurofeedback group significantly improved their relative error rate reduction (RERR) from pre-to post-training, as opposed to controls (Fig. A.2a). Both groups improved in terms of behavioral sensitivity performance, A’ (Fig. A.2b), but insignificantly so. There was no significant difference between groups. Hence, there is limited evidence whether neurofeedback *per se* provided lasting effects in sustained attention abilities.

## 4. Conclusions and Perspectives

In this study, we presented a Python-based closed-loop neurofeedback software framework for EEG. As a proof of concept, we validated our framework using a consumer-grade, dry-electrode EEG system and demonstrated real-time neuroimaging, cognitive state classification and feedback in a wearable setting. The proposed framework provides an easily extendable platform for development of real-time EEG experimental paradigms and clinically applicable tools.

### 4.1. Challenges and prospects for neurofeedback systems

The following section consolidates our contribution to the EEG neurofeedback field and highlights areas for optimization, structured as four issues that were identified by [2] as the main future challenges for visual attention-based neurofeedback systems.

1. **Filtering out noise:** Physiological, mechanical and electrical noise artifacts is a major issue in all EEG-based systems. Artifacts, of which eye artifacts are most common, can be partly solved by interstimulus intervals (ISI) for blinking. Besides implementation of ISI, artifacts are usually handled by rejection or correction of EEG epochs with e.g. suspiciously high signal amplitude values. However, both ISI and rejection of epochs reduce the ecological validity of neurofeedback systems as delay is imposed (for reviews, see [70, 109, 110]). We propose a decomposition-based artifact rejection method, SSP, in which noise components are computed throughout the experiment and automatically rejected for all EEG epochs. Specifically, through decomposition of the EEG signal, components explaining more than 10% variance were identified and discarded from the signal. The EEG components were projected back, thereby reconstructing an artifact-free signal. The components were re-estimated continuously throughout the session to ensure an updated estimate of the signal composition. In this manner, the computed noise components were participant-specific and were allowed to change throughout the time course of the neurofeedback session. For some participants, the noise components consisted of loose EEG electrodes, while for other participants it consisted of muscle tension. These components were effectively identified for each participant and improved the ability to decode the signal of interest. The technique is however not perfect, and as previously mentioned, relevant brain signal might have been filtered out and some noise signals might still be present. Machine learning approaches for further automation of these decomposition-based rejection methods have been proposed [111, 112]. Another approach to minimize the effect of artifacts is by proper feature selection representing the signal of interest. However, the challenge is to perform this feature selection in a completely automated and not computationally expensive manner. The goal of neurofeedback training is often uptraining of specific spectral bands (e.g. [72, 73, 74, 75, 76]), hence diminishing the need for feature extraction besides identification of the spectral component of interest. This approach is not optimal for neurofeedback paradigms that aim to decode more complex cognitive states, as it diminishes the possibility of using potentially relevant neural processing taking place in spectral bands other than the preselected band. In offline EEG studies, the use of specific ERP components for single-trial decoding have been demonstrated with success (e.g. [78, 85, 106]), but there is often a manual pre-selection taking place. More automated approaches include e.g. iterative variable selection based on spectral and ERP features for each participant [113] or neural networks for extraction of EEG features with subsequent classification [114, 115]. Generally, these methods are computationally expensive, but hold great potential for optimization of feature extraction in real-time neurofeedback systems. Lastly, it is important to note that if automated feature selection is not performed properly, selection and hence training of unwanted brain functions might occur - particularly relevant in clinical neurofeedback training to avoid enhancing individual‘s disabilities [2].
2. **Reliable criteria to quantify neurofeedback training effects:** There is no consensus on how to define success in a neurofeedback training study. Cognitive improvement of e.g. attention is challenging to quantify by behavioral tests due to factors unrelated to training: Inter-individual abilities in learning capacity, general ability to concentrate, motivation, mood, and personality (see review by [116]). Moreover, certain individuals fail to regulate brain activity even after repeated training session, denoted as “BCI illiterate”. This has been suggested to apply to 20% of individuals [117] or even up to 37% depending on the success threshold [118]. We conducted a pre- and post-training session for quantification of neurofeedback training outcome. Since these sessions were conducted on different days, the performance will naturally differ due to factors innate to attentional improvement, possibly outweighing the subtle training effects of training. Optimally, these factors would be controlled for, or new and more robust evaluation metrics need to be implemented. Lastly, besides a standardization of evaluation metrics, recent reviews [4, 29, 51, 50] call for homogeneity of double-blinded experimental procedures and number of training sessions. While deBettencourt and colleagues report a significant training effect after a single fMRI neurofeedback session using the implemented paradigm [60], the number of neurofeedback sessions generally range from five to twelve training sessions to produce lasting training effects [72, 45, 119, 120, 76]. We report no significant difference between neurofeedback participants and control participants after a single EEG neurofeedback training session.
3. **Accounting for intra- and inter-individual variability:** The use of neurofeedback systems outside laboratories depends on robustness to (i) Changing responses to identical conditions within individuals, and (ii) Diverse mental representations across individuals. Effort is devoted to the understanding of intra-individual and intra-study variability in BCI applications (for reviews, see [121, 116]). In the current study, the real-time decoding error rates were obtained across a wide range of EEG responses (Fig. A.1). Our system demonstrated a high inter-individual robustness and the possibility of adapting to a wide range of mental representations of visual attention across participants in an automated manner. However, the intra-individual variability was less robust: Participants who displayed highly variable responses to the same stimuli conditions (evidenced by large confidence intervals in Fig. A.1) displayed higher decoding error rates. The issue of high intra-individual variability was partly solved by two aspects of the proposed framework: (i) Implementation of a weighted moving average of EEG epochs, and (ii) Inclusion of the most recent ‘feedback’ blocks in the classifier training set (as opposed to solely using ‘stable’ blocks). Since the EEG signals during ‘feedback’ blocks were influenced by a non-stationarity of feedback and task difficulty, the inclusion of these blocks increased system robustness by training a classifier on a more dynamic set of brain responses. Lastly, our ability to decode attentional states would most likely have improved if the classifier had been updated more frequently throughout the neurofeedback session.
4. **Individual use:** Ordikhani-Seyedlar et al. highlight the restriction of BCI usage by dependence on experts for EEG set-up, running training protocols and system maintenance [2]. We highly prioritized engineering an automated, adaptable system for individual use in three different ways. First, we used a consumer-grade, user-friendly EEG system with dry electrodes. The portability and simple system set-up provide an opportunity of studying the brain in natural settings and for applications in real-life scenarios. Second, we engineered a system without the need for any manual control before or throughout the neurofeedback session. The system can easily be set up by the user with no prior technical EEG experience, thus increasing the ecological validity and usage potential of our system. Third, our system does not require any EEG recordings prior to the neurofeedback session. The system uses the first approximately 10 minutes of recorded EEG data for training a participant-specific classifier, which is used and updated throughout the session, diminishing the need for prior EEG recordings, calibration or customization.

## 5. Code and Data Availability

The code as well as sample data for the proposed neurofeedback framework is available here: https://github.com/gretatuckute/ClosedLoop/.

In accordance with our participant consent form, data is available for research purposes upon request.

## 6. Supplementary Video

A supplementary video of the neurofeedback system is available here: https://www.youtube.com/watch?v=Ns8AHIg_Wtc&feature=youtu.be

## 8. Acknowledgements

This work was supported by the Novo Nordisk Foundation Interdisciplinary Synergy Program 2014 (Biophysically Adjusted State-Informed Cortex Stimulation (BASICS)) (NNF14OC0011413).

We would like to thank Ivana Konvalinka, Per Bækgaard and Tobias Andersen at DTU Cognitive Systems for feedback on the experimental and technical aspects of the neurofeedback system.

## Appendix A.

**Figure A.1:**
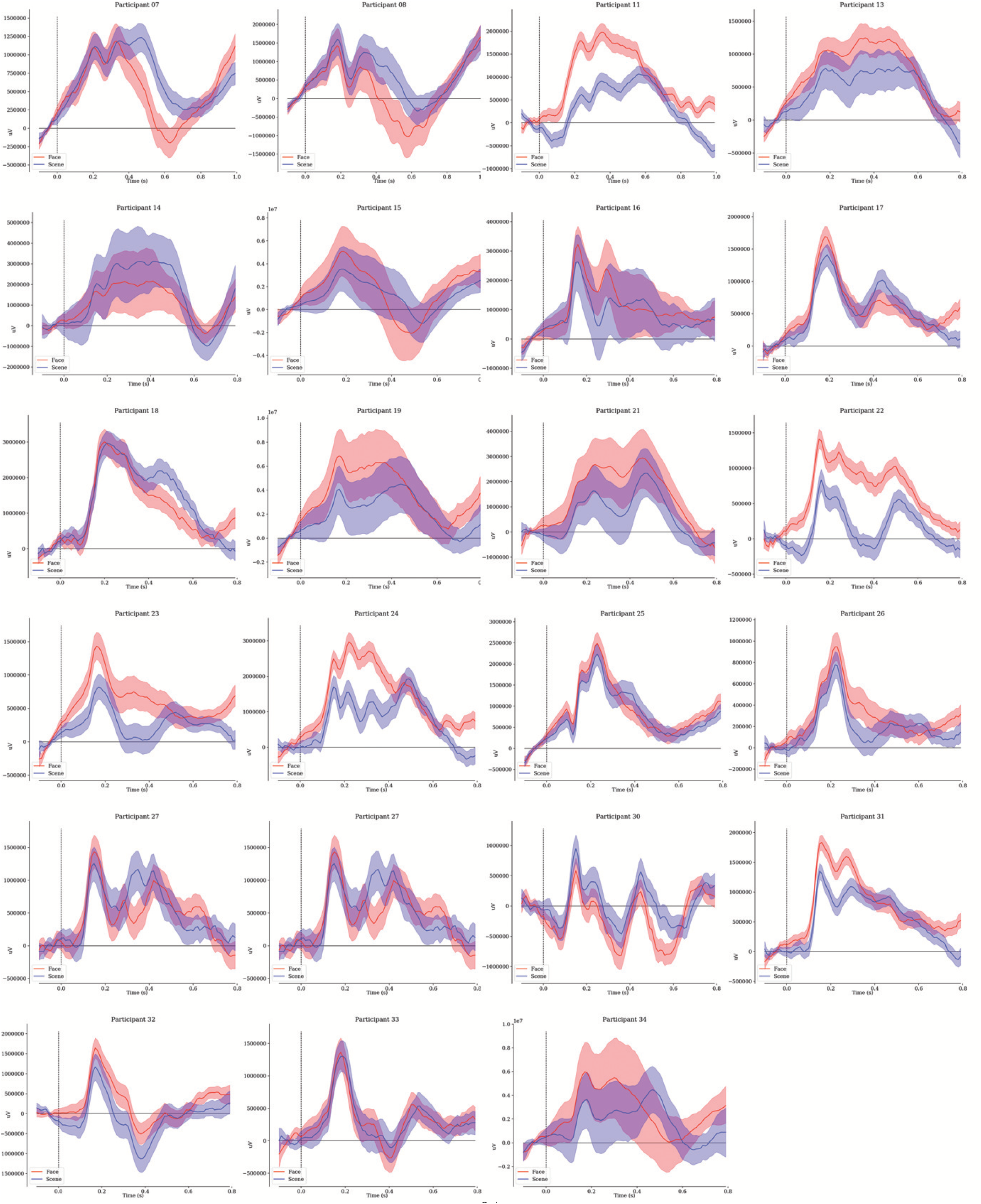
Meaned ERPs of attentional states towards faces (red) versus scenes (blue) for individual participants for occipital electrodes O1, O2, Oz, PO3 and PO4. The ERPs are extracted solely from stable blocks. Shaded regions depict 95% CI.

**Figure A.2:**
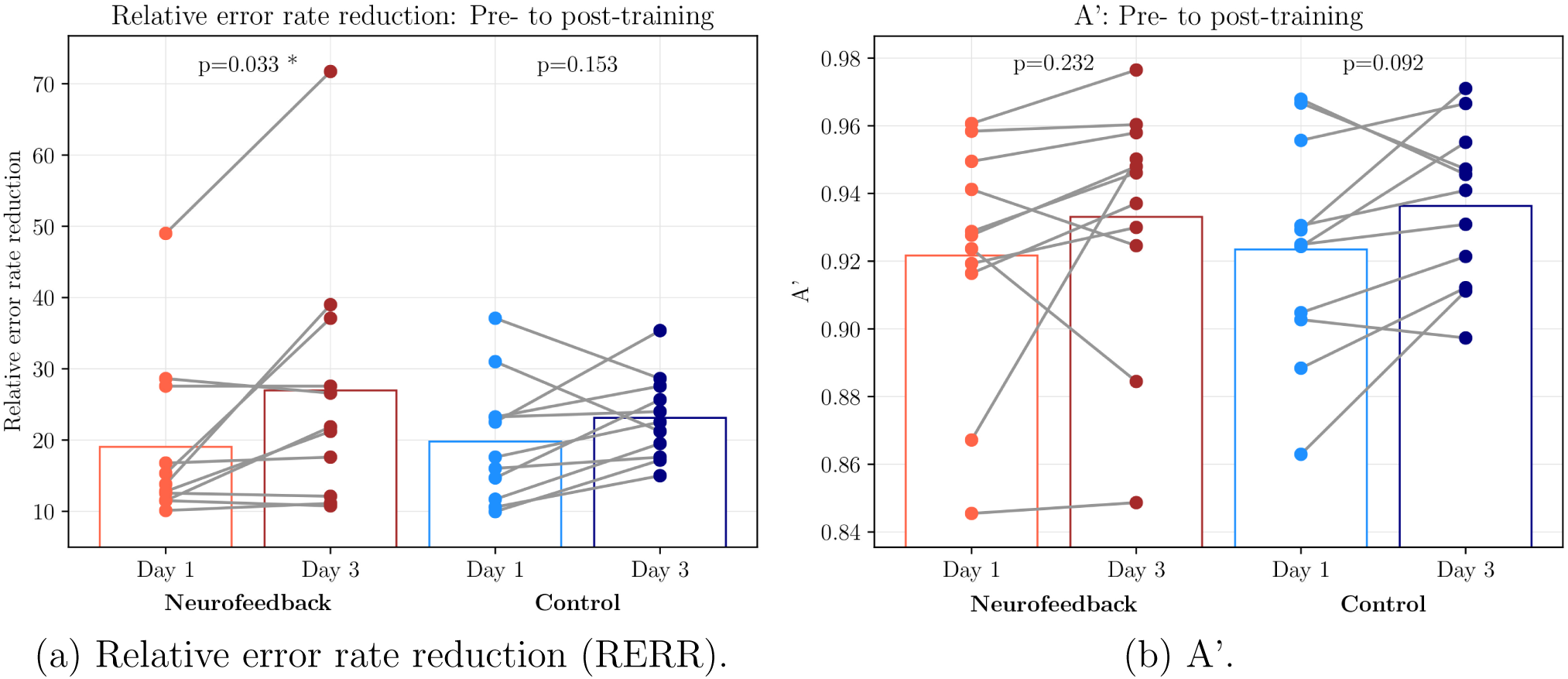
Change in behavioral performance from pre- to post-training session (day 1 to 3). Bars indicate the group-wise mean. Grey lines denote the within-participant change. Note: There was no significant effect between groups.

## Footnotes

1 www.bci2000.org

2 www.openvibe.inria.fr

3 www.neuroimage.usc.edu/brainstorm/

4 www.sccn.ucsd.edu/wiki/BCILAB

5 www.fieldtriptoolbox.org

6 www.github.com/nikolaims/nfb

7 www.brainmaster.com

8 www.myndlift.com

9 www.thoughttechnology.com

10 www.beemedic.com

11 www.neuropype.io

12 www.github.com/sccn/labstreaminglayer

13 www.nist.gov/itl/iad/image-group/color-feret-database

14 www.groups.csail.mit.edu/vision/SUN

